# Modified dN/dS for accounting transition and transversion frequency difference and non-sense substitution in genomes

**DOI:** 10.1101/2022.01.22.477328

**Authors:** Ruksana Aziz, Piyali Sen, Pratyush Kumar Beura, Debashis Das, Madhusmita Dash, Nima Dondu Namsa, Ramesh Chandra Deka, Edward J Feil, Siddhartha Sankar Satapathy, Suvendra Kumar Ray

**Author notes:** Correspondence: S. S. Satapathy, S. K. Ray.

## Abstract

The dN/dS value is estimated in homologous protein coding gene sequences between two closely related organisms for studying selection on the genes. In the usual method of calculation of synonymous (S) and non-synonymous (NS) sites in codons, the transition and transversion rates are considered same as well as no difference of pretermination codons from the other codons regarding NS substitutions is considered. In this study we are proposing a modification in the method by estimating the S and the NS sites in codons by considering difference between the transition and transversion rates and the NS substitutions leading to non-sense codons in pretermination codons. So, the dN/dS value calculated by our approach was higher than that calculated by the earlier method. The modified method was applied in estimating dN/dS in 29 homologous gene sequences of *Escherichia coli* and *Salmonella enterica*. Impact of codon degeneracy and pretermination codons on the dN/dS values estimated by our method were observed clearly. Our method of estimation that considers the above features is a realistic representation of dN/dS values in coding sequences.

## Introduction

It is usual in molecular evolution that comparison between two homologous protein coding gene sequences of two closely related species or strains is done to find out selection on the gene sequence by calculating dN/dS (Yang 1998; Hurst 2002), where dN is defined as the number of non-synonymous changes per non-synonymous site in the gene sequences whereas dS is defined by the number of synonymous changes per synonymous sites in the gene (Yang et al. 2000). Comparing the given pairs of codons in a sequence, synonymous sites at each of the three positions in a codon is calculated by finding out the fraction of the three possible substitutions at any site is being synonymous followed by summing up the synonymous sites at the three positions (Yang and Nielsen 2002). Similarly, non-synonymous sites at each of the three positions in a codon is calculated by finding out the fraction of the three changes is being non-synonymous followed by summing up the non-synonymous sites at the three positions (Nei and Gojobori, 1986). The method has been widely used by many researchers and many critical reviews have been made on its application after the proposition by Gojobori and Nei in 1986 (Hurst 2002; Rocha et al 2006; Kryazhimskiy and Plotkin, 2008; Weber et al 2014; Spielman and Wilke, 2015). Using computer simulation detail analysis of estimating S and NS sites in codons proposed by different researchers have been studied earlier (Ina 1995).

At every position of a codon, out of the three substitutions, one is a transition (*ti*) while the other two are transversions (*tv*) (Gojobori et al. 1982; Lyons and Lauring 2017). It is already known that *ti* is more frequent than *tv* (Gojobori *et al*. 1982; Petrov and Hartl 1999; Sen *et al* 2021). In *E. coli*, a *ti* substitution is in average four times more frequent than a *tv* substitution (Sen et al., 2021). Therefore, it is obvious that if a codon is only undergoing synonymous substitutions due to *ti*, as observed in case of a codon with two-fold degenerate site, the relative synonymous frequency is going to be higher than a codon with four-fold degenerate site, where the codon is undergoing synonymous substitutions due to both *ti* and *tv*. So synonymous and non-synonymous site calculation for a *ti* and for a *tv* should be considered differently. It is pertinent to consider the difference between *ti* and *tv* rates while calculating the synonymous sites (S) and non-synonymous (NS) sites for a codon (Gojobori et al. 2015).

Apart from the *ti* and *tv* rate difference, we have to also consider the difference between pretermination codon from the other codons while calculating the NS site of the codons (Supplementary Table 1). Unlike other non-synonymous changes, purifying selection against the nonsense codon is maximum. Therefore, substitution frequency in pretermination codons is likely to be observed lesser than the other codons. This might be an explanation for the observation of lower substitution in UGG (Trp) codon in comparison to the AUG (Met) codon (Andersson and Kurland 1991). Though, low frequency of non-synonymous substitutions in Trp and Cys codons have been attributed of the possible positional significance in protein function (Heizer Jr et al. 2006), our explanation can also be an additional one. In our recent study in *E. coli* gene sequence comparison, we have observed that substitutions in Trp and Met codons are comparable when the non-sense codons will not be considered (unpublished work). Therefore, while calculating the non-synonymous site in case of pretermination codons, number of changes resulting nonsense codon might not to be considered. Similar explanations can be given for other pretermination codons. In case of UUA and UUG, the NS sites are different (Table 1.2) despite both being pretermination codons because in case of UUA two different substitutions results nonsense codons while in case UUG only one substitution results in a nonsense codon.

**Table 1.1:** S and NS sites of codons in the genetic code table by the earlier method.

| Old calculation |  |  |  |  |  |  |  |  |  |
| --- | --- | --- | --- | --- | --- | --- | --- | --- | --- |
|  | U |  | C |  | A |  | G |  |  |
|  | S | NS | S | NS | S | NS | S | NS |  |
| U | 0.333 | 2.667 | 1.000 | 2.000 | 0.333 | 2.667 | 0.333 | 2.667 | U |
|  | 0.333 | 2.667 | 1.000 | 2.000 | 0.333 | 2.667 | 0.333 | 2.667 | C |
|  | 0.667 | 2.333 | 1.000 | 2.000 | N/A | N/A | N/A | N/A | A |
|  | 0.667 | 2.333 | 1.000 | 2.000 | N/A | N/A | 0.000 | 3.000 | G |
| C | 1.000 | 2.000 | 1.000 | 2.000 | 0.333 | 2.667 | 1.000 | 2.000 | U |
|  | 1.000 | 2.000 | 1.000 | 2.000 | 0.333 | 2.667 | 1.000 | 2.000 | C |
|  | 1.333 | 1.667 | 1.000 | 2.000 | 0.333 | 2.667 | 1.333 | 1.667 | A |
|  | 1.333 | 1.667 | 1.000 | 2.000 | 0.333 | 2.667 | 1.333 | 1.667 | G |
| A | 0.667 | 2.333 | 1.000 | 2.000 | 0.333 | 2.667 | 0.333 | 2.667 | U |
|  | 0.667 | 2.333 | 1.000 | 2.000 | 0.333 | 2.667 | 0.333 | 2.667 | C |
|  | 0.667 | 2.333 | 1.000 | 2.000 | 0.333 | 2.667 | 0.666 | 2.334 | A |
|  | 0.000 | 3.000 | 1.000 | 2.000 | 0.333 | 2.667 | 0.666 | 2.334 | G |
| G | 1.000 | 2.000 | 1.000 | 2.000 | 0.333 | 2.667 | 1.000 | 2.000 | U |
|  | 1.000 | 2.000 | 1.000 | 2.000 | 0.333 | 2.667 | 1.000 | 2.000 | C |
|  | 1.000 | 2.000 | 1.000 | 2.000 | 0.333 | 2.667 | 1.000 | 2.000 | A |
|  | 1.000 | 2.000 | 1.000 | 2.000 | 0.333 | 2.667 | 1.000 | 2.000 | G |

**Table 1.2:** S and NS sites of codons in the genetic code table by the new proposed method.

| New calculation |  |  |  |  |  |  |  |  |  |
| --- | --- | --- | --- | --- | --- | --- | --- | --- | --- |
|  | U |  | C |  | A |  | G |  |  |
|  | S | NS | S | NS | S | NS | S | NS |  |
| U | 0.667 | 2.333 | 1.000 | 2.000 | 0.667 | 2.000 | 0.667 | 2.167 | U |
|  | 0.667 | 2.333 | 1.000 | 2.000 | 0.667 | 2.000 | 0.667 | 2.167 | C |
|  | 1.333 | 1.333 | 1.000 | 1.667 | N/A | N/A | N/A | N/A | A |
|  | 1.333 | 1.500 | 1.000 | 1.833 | N/A | N/A | 0.000 | 1.667 | G |
| C | 1.000 | 2.000 | 1.000 | 2.000 | 0.667 | 2.333 | 1.000 | 2.000 | U |
|  | 1.000 | 2.000 | 1.000 | 2.000 | 0.667 | 2.333 | 1.000 | 2.000 | C |
|  | 1.667 | 1.333 | 1.000 | 2.000 | 0.667 | 1.667 | 1.167 | 1.167 | A |
|  | 1.667 | 1.333 | 1.000 | 2.000 | 0.667 | 1.667 | 1.167 | 1.833 | G |
| A | 0.833 | 2.167 | 1.000 | 2.000 | 0.667 | 2.333 | 0.667 | 2.333 | U |
|  | 0.833 | 2.167 | 1.000 | 2.000 | 0.667 | 2.333 | 0.667 | 2.333 | C |
|  | 0.333 | 2.667 | 1.000 | 2.000 | 0.667 | 2.167 | 0.833 | 2.000 | A |
|  | 0.000 | 3.000 | 1.000 | 2.000 | 0.667 | 2.167 | 0.833 | 2.167 | G |
| G | 1.000 | 2.000 | 1.000 | 2.000 | 0.667 | 2.333 | 1.000 | 2.000 | U |
|  | 1.000 | 2.000 | 1.000 | 2.000 | 0.667 | 2.333 | 1.000 | 2.000 | C |
|  | 1.000 | 2.000 | 1.000 | 2.000 | 0.667 | 2.167 | 1.000 | 1.833 | A |
|  | 1.000 | 2.000 | 1.000 | 2.000 | 0.667 | 2.167 | 1.000 | 2.000 | G |

## Materials and Methods

Calculation of S and NS in the modified method

### Existing Method

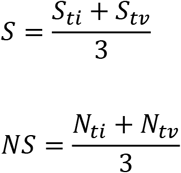

### Proposed Method

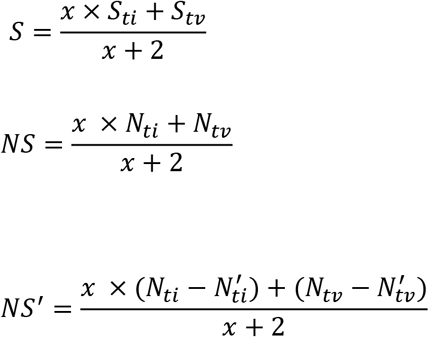

*S*: synonymous site of a codon
*NS*: non-synonymous site of a codon
*x* = number of times a *ti* is more frequent than a *tv*
*S_ti_* = Synonymous transition due to single site substitution in a codon
S_tv_= Synonymous transversion due to single site substitution in a codon
*N_ti_*= Non-synonymous transition due to single site substitution in a codon
*N_tv_*= Non-synonymous Transversion due to single site substitution in a codon
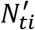 = Number of *N_ti_* resulting to stop codon in a codon
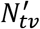= Number of *N_tv_* resulting to stop codon in a codon

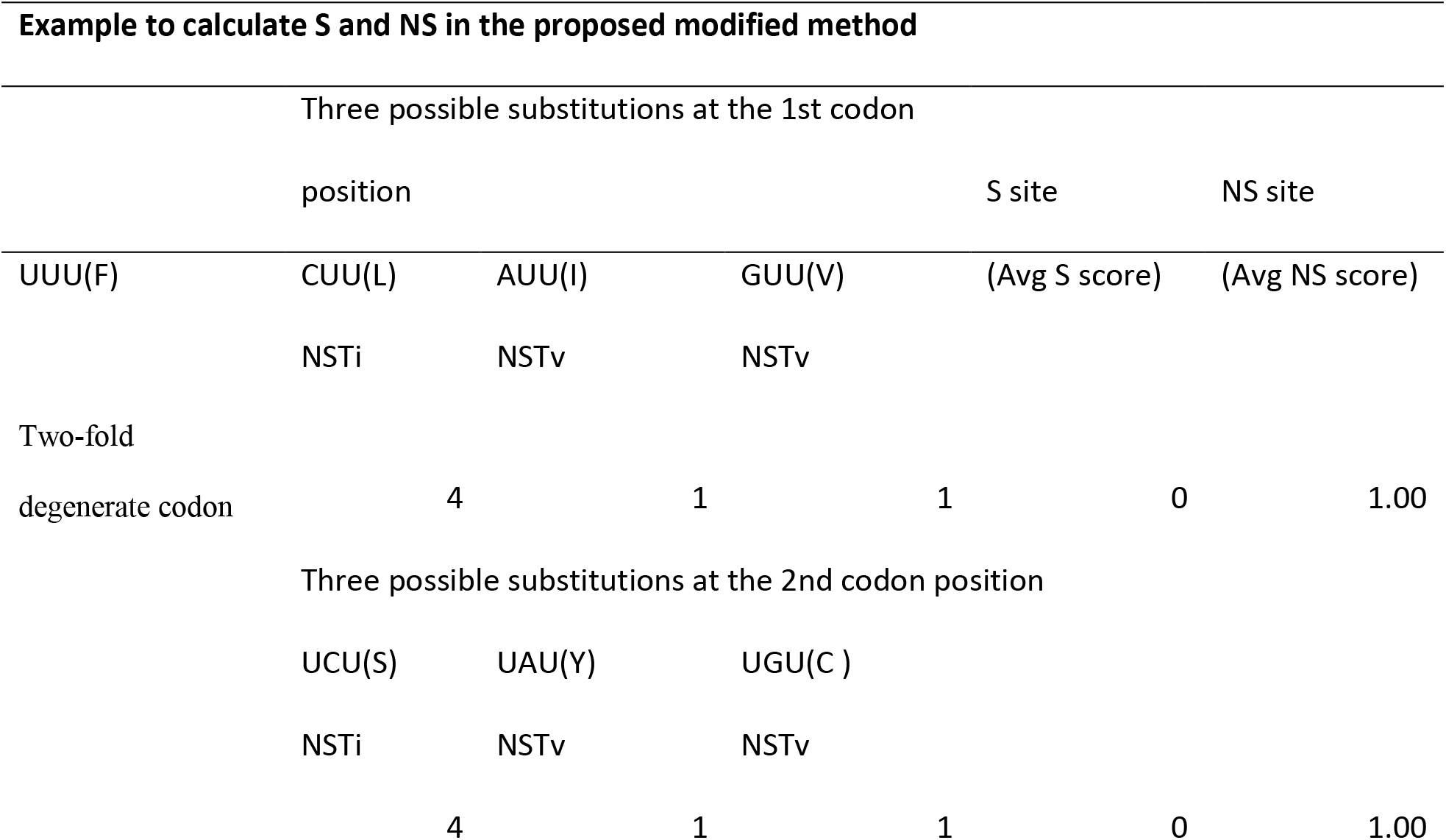
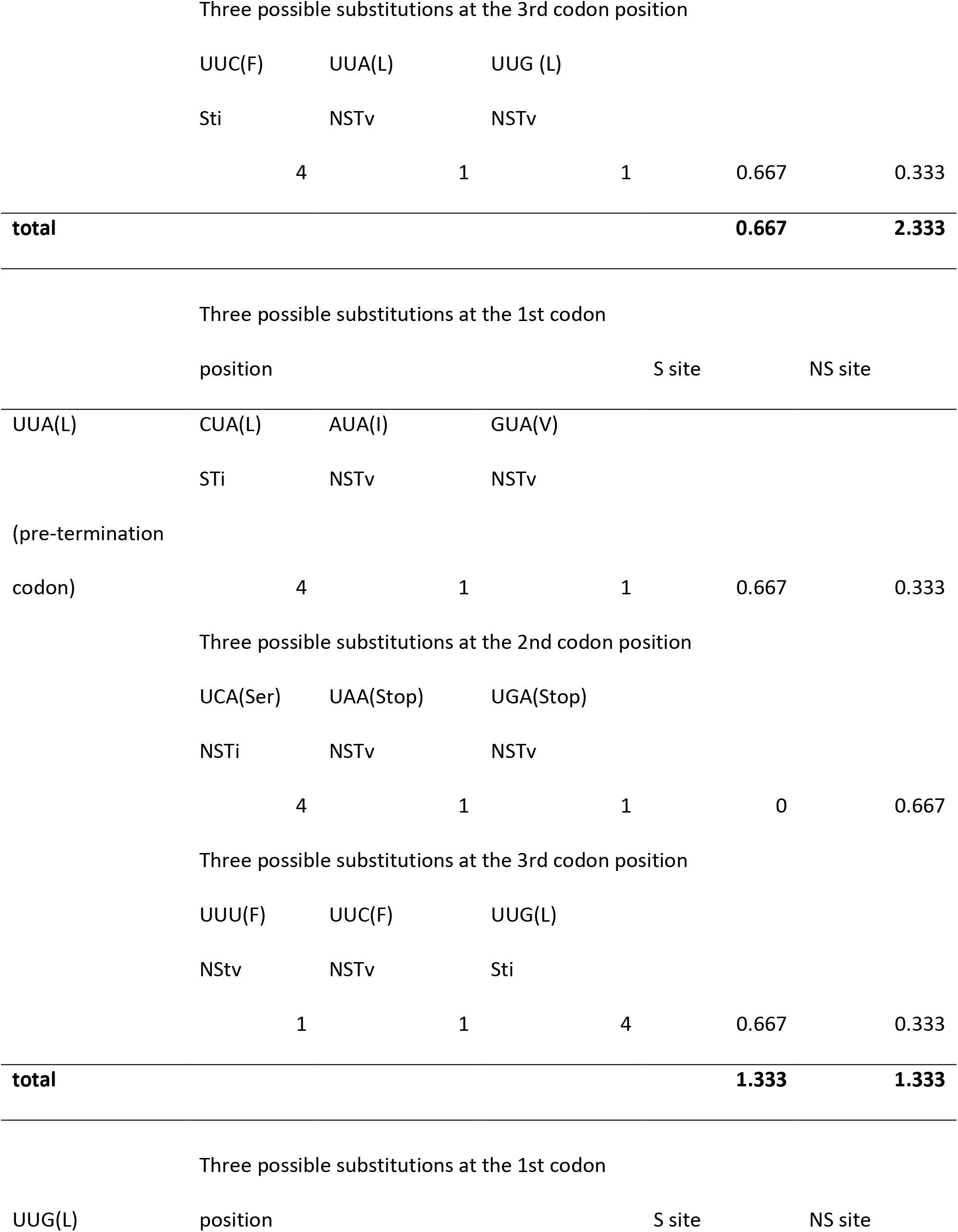
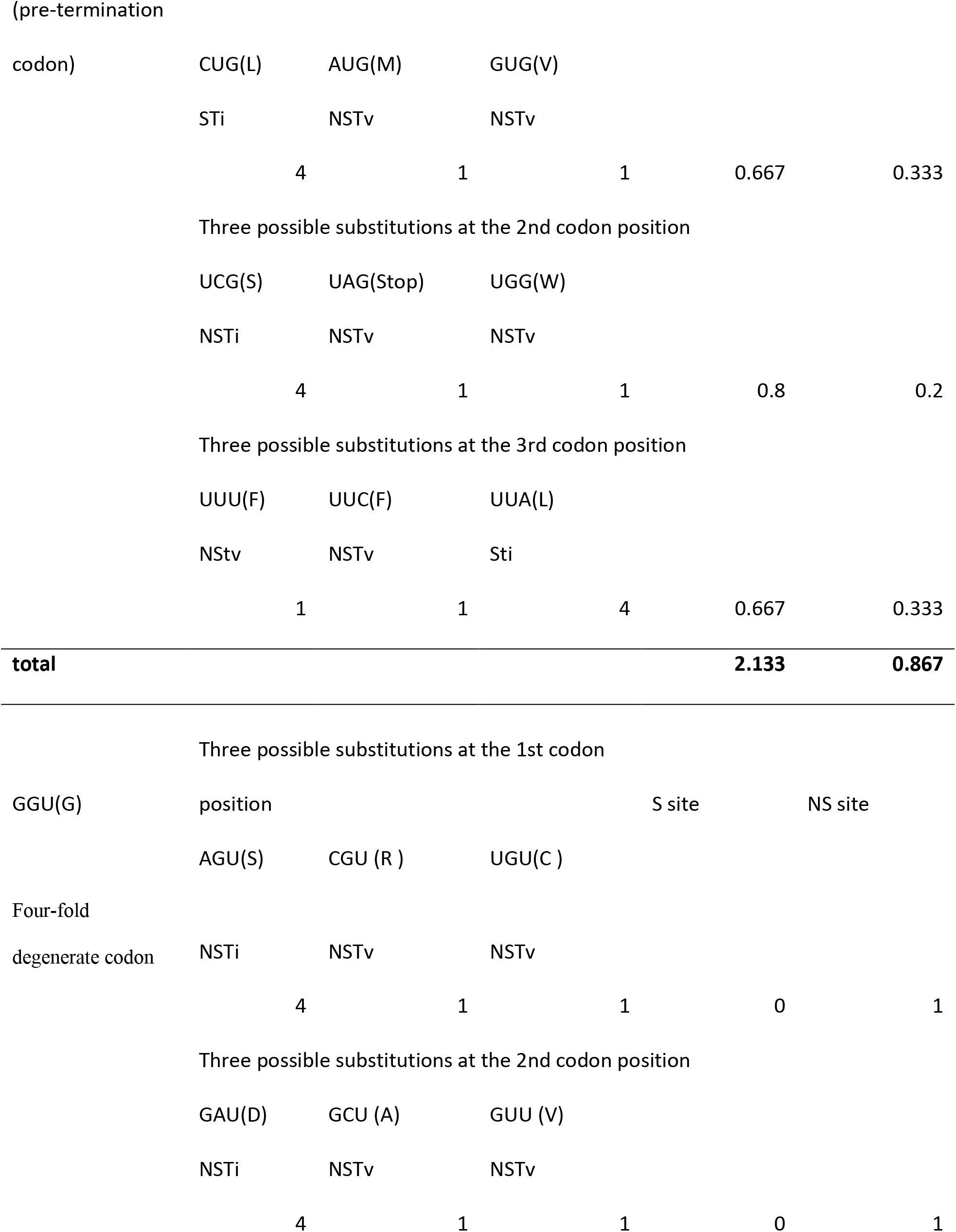
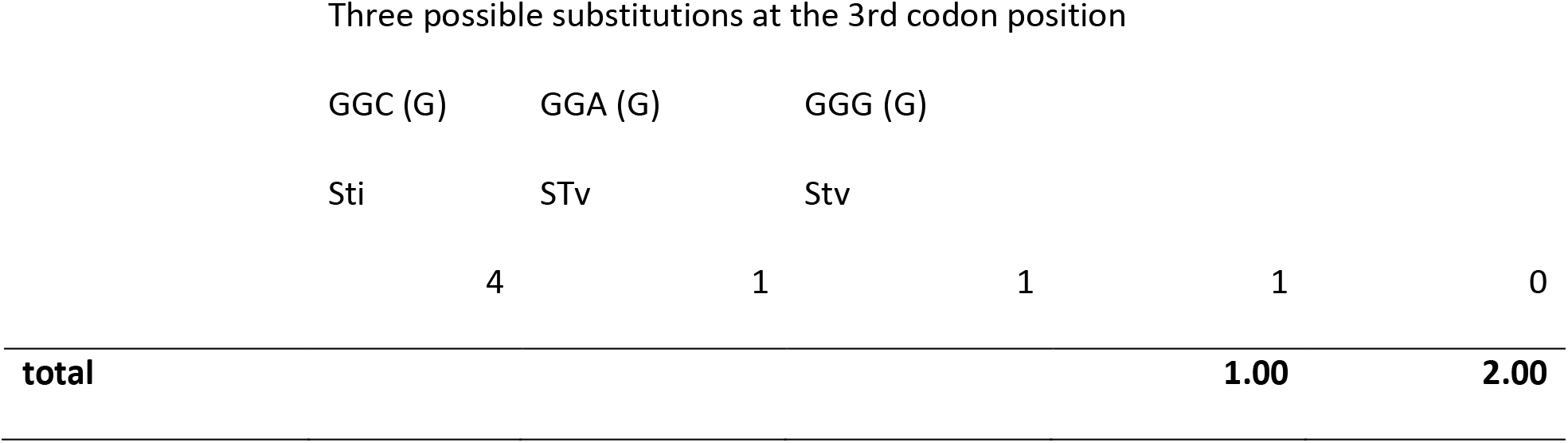

## Results and Discussion

The equation for the calculation S and NS sites of a codon by both the old method and the new method is given in the Materials and Methoids. Here, we are presenting a genetic code table with the S and NS site values of codons used by the researchers to calculate dN/dS values and another table with the proposed modified S and NS site values of codons calculated considering *ti* and *tv* rate difference proposed by us as well as considering the non-synonymous substitutions leading to non-sense codons in case of pretermination codons. In comparison to the S and NS site values of codons in the earlier method given in the genetic code table (Table 1.1), we can observe that there are differences of the S and NS site values of codons in the new method given in the genetic code table (Table 1.2). Some of the examples are given as follows. The S site for CUG/CUA codons is 1.333, which is same as for CGA/CGG codons in the earlier method. In the proposed method, the S site for CUA/CUG codons is 1.667, and that for the CGA/CGG codons is 1.166. It is known that synonymous substitution in CUA/CUG codon is the highest among all the codons in the genetic code table because it involves two transition and two transversion substitutions while synonymous substitution in case of CGA/CGG codon involves one transition and three transversion substitution (Dasso and Jackson 1989). Further, the S site value of AUA codon in the earlier table is 0.667 whereas the S site value of the codon in the proposed table is 0.333, which is the least among the codons that can go synonymous substitutions in the genetic code table. Therefore, the values in the proposed table can be an easy reference for understanding the rate of synonymous substitutions expected in different codons, that is based on difference between *ti* and *tv* rates (Osawa et al. 1989; Yang and Yoder 1999). It is pertinent to note that this approach is not limited to only when *ti* is four times more than *tv* considered here for *E. coli* (Table 1.2) but is suitable for any fold increase of the *ti* values than *tv* (Supplementary Table 1). Regarding non-synonymous substitutions, the obvious observation is between AUG (Met) and UGG (Trp) codons. UGG is a pretermination codon but AUG is not. In case of UGG, two (transitions) out of nine substitutions result non-sense codon. In the earlier table (Table 1.1) the NS site values are considered as 3.0 for both, which is suggesting that both the codons are equal regarding NS substitutions. However, in the proposed modified table (Table 1.2) in this study, the NS site value for AUG is considered as 3.0 while for UGG is considered as 1.667.

It is observed in the earlier and modified Tables that the value of synonymous sites is 53.333 (~30 %) in the modified method while the same in the earlier method is 44.664 (~25 %) indicating the increase in the synonymous sites by the new calculation as it is known that higher proportion of synonymous changes attributes to transitions. Further, sum of all the non-synonymous sites value is 123.335 (~70 %) in the modified calculation while the same is 138.336 (~75%) in the earlier method, indicating the decrease in the non-synonymous sites due to consideration of non-sense sites in the pretermination codons by the new calculation. Therefore, the estimated dN/dS values by our method for genes are likely to be higher than the previous calculations.

A computer programme is developed considering the proposed modification in calculating the S and NS sites as well as the new dN/dS values. We then tested our method in a sample set of 29 genes of two closely related Gram-negative bacteria i.e. *E. coli* and *S. enterica*. These genes are of same length in both the bacteria having no insertion or deletion mutations. We calculated dN/dS both by the previous approach as well as by the proposed new approach (Table 2). It is evident from the result that the dN/dS values using the new calculation is higher than that calculated by the previous method in each of the 29 genes. This is in concordance with our proposition made above. Further we found out percentage increase in the dN/dS values in each gene (Table 2). Among these genes, the increase in dN/dS values range from 32 to 50%. To explore the reason for this difference by finding out the number of two-fold degenerate (TFD) codons vs number of four-fold degenerate (FFD) codons (McClellan 2000). If the proportion of TFD is more than the FFD, then the difference values are likely to be more. We found out the ratio of TFD and FFD in each of the 29 genes. The minimum ratio is 0.680:1.000 while the maximum ratio was 1.718:1.000. We did a correlation between the difference values (Table 2) with the TFD:FFD ratio. The pearson *r* is observed as 0.933 of the correlation between difference values vs the ratio of TFD:FFD codons in each gene. This proved our approach is considering the degeneracy of the codons unlike the earlier method of calculation. We also studied the impact of pretermination codon composition in genes and the dN/dS values. Composition of pretermination codons varies from 16.0 to 36.0 % across these 29 genes (Table 2). We then did a correlation between pretermination codon % in a gene and the % increase in the dN/dS values calculated by the new method. The pearson *r* value is 0.640 suggesting the impact of pretermination codon composition in a gene on the dN/dS value. This also proved our method of calculation of dN/dS is influenced by the composition of pretermination codon.

**Table 2:** A comparison of dN/dS values by the old and the new methods in 29 genes of *Escherichia coli* and *Salmonella enterica*.

| Sl | Gene | Number<br>of S<br>Mutation | Number<br>of NS<br>Mutation | S site<br>(New) | NS site<br>(New) | dN/dS<br>(New) | S site<br>(Old) | NS site<br>(Old) | dN/dS<br>(Old) | Diff | %<br>increase | Ratio | Freq<br>PreTer |
| --- | --- | --- | --- | --- | --- | --- | --- | --- | --- | --- | --- | --- | --- |
| 1 | araE | 223 | 23 | 410.42 | 966.84 | 0.04 | 340.83 | 1078.17 | 0.03 | 0.011 | 34.280 | 0.680 | 0.205 |
| 2 | gltI | 133 | 20 | 256.25 | 621.43 | 0.06 | 202.50 | 706.50 | 0.04 | 0.019 | 43.867 | 1.337 | 0.274 |
| 3 | gltR_1 | 171 | 76 | 252.75 | 591.01 | 0.19 | 209.50 | 668.00 | 0.14 | 0.051 | 36.361 | 0.846 | 0.248 |
| 4 | gltX | 171 | 30 | 398.58 | 971.35 | 0.07 | 314.67 | 1101.33 | 0.05 | 0.022 | 43.619 | 1.366 | 0.248 |
| 5 | hemA | 163 | 28 | 370.92 | 840.93 | 0.08 | 299.83 | 957.17 | 0.05 | 0.022 | 40.807 | 1.111 | 0.265 |
| 6 | hemB | 149 | 27 | 277.58 | 671.01 | 0.07 | 226.17 | 748.83 | 0.05 | 0.020 | 36.969 | 1.037 | 0.255 |
| 7 | hemC | 145 | 45 | 291.33 | 645.43 | 0.14 | 244.17 | 718.83 | 0.11 | 0.035 | 32.888 | 0.774 | 0.224 |
| 8 | hemD | 82 | 57 | 224.00 | 480.42 | 0.32 | 182.00 | 559.00 | 0.23 | 0.098 | 43.207 | 1.068 | 0.328 |
| 9 | hemG | 79 | 30 | 150.33 | 370.67 | 0.15 | 121.67 | 424.33 | 0.11 | 0.045 | 41.448 | 1.302 | 0.302 |
| 10 | hemH | 144 | 46 | 283.50 | 644.84 | 0.14 | 234.00 | 729.00 | 0.10 | 0.038 | 36.965 | 0.927 | 0.265 |
| 11 | hemN_1 | 175 | 51 | 330.42 | 762.26 | 0.13 | 262.33 | 874.67 | 0.09 | 0.039 | 44.527 | 1.246 | 0.280 |
| 12 | hemN_2 | 185 | 46 | 394.83 | 925.68 | 0.11 | 304.67 | 1069.33 | 0.07 | 0.035 | 49.707 | 1.718 | 0.288 |
| 13 | hisB | 149 | 23 | 302.67 | 732.93 | 0.06 | 238.33 | 829.67 | 0.04 | 0.019 | 43.754 | 1.408 | 0.247 |
| 14 | hisF | 102 | 18 | 220.08 | 530.68 | 0.07 | 180.67 | 596.33 | 0.05 | 0.020 | 36.888 | 1.044 | 0.232 |
| 15 | lacG | 109 | 30 | 242.92 | 578.00 | 0.12 | 202.83 | 643.17 | 0.09 | 0.029 | 33.263 | 0.680 | 0.163 |
| 16 | leuE | 119 | 45 | 192.42 | 427.34 | 0.17 | 158.33 | 480.67 | 0.12 | 0.046 | 36.692 | 0.868 | 0.249 |
| 17 | leuS | 278 | 50 | 720.42 | 1772.37 | 0.07 | 582.83 | 2000.16 | 0.05 | 0.021 | 39.491 | 1.112 | 0.259 |
| 18 | malF | 196 | 60 | 453.33 | 1037.51 | 0.13 | 366.67 | 1178.33 | 0.10 | 0.039 | 40.418 | 0.917 | 0.237 |
| 19 | pepN | 354 | 66 | 749.75 | 1779.12 | 0.08 | 595.00 | 2017.99 | 0.05 | 0.024 | 42.925 | 1.303 | 0.240 |
| 20 | polA | 440 | 74 | 817.50 | 1885.80 | 0.07 | 658.50 | 2128.50 | 0.05 | 0.021 | 40.124 | 1.086 | 0.279 |
| 21 | recO | 109 | 23 | 219.42 | 486.51 | 0.10 | 178.83 | 550.17 | 0.07 | 0.027 | 38.747 | 0.975 | 0.247 |
| 22 | recR | 73 | 12 | 177.33 | 408.01 | 0.07 | 146.67 | 459.33 | 0.05 | 0.019 | 36.121 | 0.971 | 0.257 |
| 23 | rpoC | 216 | 27 | 1265.75 | 2858.56 | 0.06 | 1041.00 | 3183.01 | 0.04 | 0.014 | 35.391 | 0.996 | 0.223 |
| 24 | rpoE | 37 | 1 | 166.33 | 386.42 | 0.01 | 134.17 | 441.83 | 0.01 | 0.003 | 41.757 | 1.175 | 0.302 |
| 25 | rpoH | 79 | 11 | 242.08 | 578.93 | 0.06 | 192.17 | 662.83 | 0.04 | 0.018 | 44.233 | 1.397 | 0.260 |
| 26 | rpoN | 186 | 31 | 413.08 | 963.35 | 0.07 | 324.67 | 1109.33 | 0.05 | 0.023 | 46.513 | 1.411 | 0.291 |
| 27 | topA_2 | 68 | 36 | 153.50 | 366.92 | 0.22 | 122.00 | 421.00 | 0.15 | 0.068 | 44.362 | 1.379 | 0.365 |
| 28 | trpA | 138 | 56 | 241.83 | 541.93 | 0.18 | 199.67 | 607.33 | 0.13 | 0.048 | 35.736 | 0.865 | 0.249 |
| 29 | trpS | 108 | 14 | 288.83 | 681.35 | 0.05 | 227.17 | 777.83 | 0.04 | 0.017 | 45.153 | 1.308 | 0.272 |
| Min |  | 37 | 1 | 150.33 | 366.92 | 0.01 | 121.67 | 421.00 | 0.01 | 0.003 | 32.888 | 0.680 | 0.163 |
| Max |  | 440 | 76 | 1265.75 | 2858.56 | 0.32 | 1041.00 | 3183.01 | 0.23 | 0.098 | 49.707 | 1.718 | 0.365 |

Ratio (TFD:FFD): Two fold degenerate site (TFD); Four-fold degenerate site (FFD)
Freq PreTer: Fraction of pre-termination codons in a gene
List of eighteen pretermination codons: UUA, UUG, UCA, UCG, UAU, UAC, UGU, UGC, UGG, CAA, CAG, AAA, AAG, GAA, GAG, CGA, AGA, GGA

We would like to add here that only the average difference between *ti* and *tv* rates has been considered to estimate the new S and NS values in this study. Consideration of individual substitution rates make the estimation more complex. Lastly, we have estimated dN/dS by finding out S sites and NS sites for codons in the genetic code table by implementing the average difference in the rate of *ti* and *tv* in organisms and the difference between pretermination and other codons. This work sheds light on the importance of the composition of amino acids with different codon degeneracy as well as pretermination nature of codons in calculation of the dN/dS values.

## Acknowledgements

RA is thankful for the JRF fellowship from the DBT grant (BT/511/NE/TBP/2013) and (BT/403/NE/U-Excel/2013). PS is thankful to UGC, GoI New Delhi for the JRF. PKB is thankful to Tezpur University for the Institutional fellowship. SSS and SKR are thankful to DBT, GoI for the twinning grant (BT/PR16361/NER/95/192/2015 date 18-10-2016) to them. SKR and RCD are thankful to DBT, GoI for the twinning grant BT/PR16182/NER/95/92/2015. SSS is thankful to DBT for the NE Overseas Associateship, which helped him to work in University of Bath. EF, SSS, RCD and SKR are thankful to the society for Molecular Biology and Evolution (SMBE) for holding the satellite meeting at Kaziranga, Assam, India on Dec 14^th^-16^th^, 2017, which helped the authors to have collaboration on this work. NDN, RCD, SKR, SSS thankfully acknowledge the DBT funded Bioinformatics and Computational Biology Centre at Tezpur University.

## Contributions of authors

RA: executed, data analysis, writing, discussion; PS: executed, data analysis, writing; PKB: executed, writing, discussion; DD, MD: computer programming, execution; RCD, NDN, EF: critical analysis, writing, discussion; SSS, SKR: design, critical analysis, writing, discussion.

All the authors endorse the manuscript

Website link for calculating dN/dS: for any queries regarding computer programme please contact SSS.

