## Supplemental Table 1 for "Modified dN/dS for accounting transition and transversion frequency difference and non-sense substitution in genomes"

**Supplementary Table 1: S and NS changes observed across 100 different strains of *E. coli* and *S. enterica***

| Organisms | | *Escherichia coli* | | | | | *Salmonella enterica* | | | | |
|---|---|---|---|---|---|---|---|---|---|---|---|
| Genes | Amino Acids | (G+C)% | S obs | NS obs | Nc | CAI | (G+C)% | S obs | NS obs | Nc | CAI |
| *araE* | 472 | 51.52 | 66 | 9 | 44.92 | 0.415 | 52.85 | 64 | 11 | 63.21 | 0.444 |
| *gltD_1* | 659 | 55.86 | 56 | 8 | 46.55 | 0.471 | 58.66 | 35 | 5 | 64.68 | 0.474 |
| *gltR_1* | 293 | 53.85 | 60 | 21 | 46.34 | 0.381 | 57.04 | 38 | 13 | 63.23 | 0.377 |
| *gltX* | 471 | 52.54 | 68 | 10 | 39.53 | 0.631 | 55.37 | 89 | 10 | 65.89 | 0.661 |
| *hemA* | 418 | 54.34 | 122 | 19 | 42.06 | 0.473 | 57.12 | 60 | 3 | 65.16 | 0.476 |
| *hemB* | 324 | 54.46 | 61 | 3 | 43.63 | 0.515 | 55.49 | 51 | 12 | 62.77 | 0.526 |
| *hemC* | 320 | 55.97 | 63 | 10 | 45.72 | 0.424 | 58.67 | 48 | 7 | 62.93 | 0.396 |
| *hemD* | 246 | 53.44 | 69 | 33 | 48.88 | 0.32 | 57.62 | 22 | 20 | 58.7 | 0.296 |
| *hemG* | 181 | 51.65 | 34 | 5 | 43.45 | 0.424 | 50.73 | 22 | 7 | 56.59 | 0.449 |
| *hemH* | 320 | 54.31 | 57 | 10 | 43.07 | 0.469 | 57.01 | 50 | 21 | 62.31 | 0.438 |
| *hemN_1* | 378 | 53.39 | 125 | 29 | 43.44 | 0.399 | 55.15 | 58 | 31 | 61.48 | 0.454 |
| *hemN_2* | 457 | 53.13 | 54 | 7 | 43.05 | 0.486 | 51.97 | 47 | 16 | 61.14 | 0.525 |
| *hisB* | 355 | 53.84 | 140 | 25 | 37.69 | 0.467 | 52.34 | 60 | 14 | 60.96 | 0.547 |
| *hisF* | 258 | 52.51 | 112 | 19 | 39.92 | 0.555 | 55.08 | 34 | 5 | 62.93 | 0.557 |
| *lacG* | 281 | 54.49 | 70 | 9 | 38.89 | 0.482 | 53.78 | 42 | 2 | 70.57 | 0.463 |
| *leuE* | 212 | 45.07 | 19 | 9 | 42.56 | 0.358 | 46.95 | 28 | 5 | 49.3 | 0.374 |
| *leuS* | 860 | 53.39 | 116 | 19 | 38.14 | 0.668 | 56.1 | 110 | 15 | 68.29 | 0.683 |
| *malF* | 514 | 53.33 | 75 | 13 | 39.47 | 0.474 | 52.69 | 78 | 13 | 65.83 | 0.476 |
| *pepN* | 870 | 54.08 | 192 | 20 | 40.78 | 0.493 | 53.73 | 106 | 29 | 62.23 | 0.558 |
| *polA* | 928 | 51.96 | 91 | 20 | 44.69 | 0.477 | 53.89 | 120 | 26 | 61.03 | 0.497 |
| *recO* | 242 | 53.91 | 44 | 5 | 42.34 | 0.408 | 55.28 | 36 | 8 | 61.73 | 0.415 |
| *recR* | 201 | 57.59 | 19 | 1 | 40.75 | 0.479 | 61.39 | 34 | 3 | 73.27 | 0.508 |
| *rpoC* | 1407 | 53.88 | 105 | 5 | 34.89 | 0.749 | 54.76 | 279 | 9 | 60.23 | 0.768 |
| *rpoE* | 191 | 48.78 | 6 | 1 | 46.72 | 0.392 | 51.04 | 10 | 1 | 54.69 | 0.357 |
| *rpoH* | 284 | 54.15 | 44 | 1 | 37.75 | 0.57 | 54.04 | 22 | 1 | 62.46 | 0.636 |
| *rpoN* | 477 | 53.14 | 59 | 9 | 46.08 | 0.432 | 54.04 | 56 | 6 | 63.18 | 0.462 |
| *topA_2* | 180 | 50.28 | 23 | 7 | 46.01 | 0.462 | 53.04 | 29 | 6 | 53.04 | 0.392 |
| *trpA* | 268 | 53.53 | 68 | 13 | 43.62 | 0.452 | 57.62 | 46 | 14 | 64.31 | 0.443 |
| *trpS* | 334 | 52.64 | 56 | 5 | 40 | 0.583 | 51.94 | 60 | 7 | 60.6 | 0.641 |

It is obvious to note that number S and NS changes among the strains within a species given in this Table in *E. coli* and in *S. enterica* is lower than the S and NS values across the two bacterial species given in Table 2.
